# Differential interactions of the proteasome inhibitor PI31 with constitutive and immuno-20S proteasomes

**DOI:** 10.1101/2023.12.11.570853

**Authors:** Jason Wang, Abbey Kjellgren, George N. DeMartino

**Author notes:** These authors contributed equally to this work (Jason Wang, Abbey Kjellgren). Correspondence should be addressed to G.N.D.

## Abstract

PI31 (<u>P</u>roteasome *I*nhibitor of <u>31</u>,000 Daltons) is a 20S proteasome binding protein originally identified as an in vitro inhibitor of 20S proteasome proteolytic activity. Recently reported cryo-electron microscopy structures of 20S-PI31 complexes reveal a surprising structural basis for proteasome inhibition. The natively disordered proline-rich C-terminus of PI31 enters the central chamber in the interior of the cylindrical 20S proteasome and interacts directly with the proteasome’s multiple catalytic threonine residues a manner predicted to inhibit their enzymatic function while evading its own proteolysis. Higher eukaryotes express an alternative form of 20S proteasome featuring genetically and functionally distinct catalytic subunits. This proteasome is expressed in tissues involved in immune function or in response to certain cytokines such as interferon-γ and has been termed “immuno-proteasome.” We examined the relative effects of PI31 on constitutive and immuno-20S proteasomes and show that PI31 inhibits the immuno-20S proteasome (20Si) to a significantly lesser degree than it inhibits constitutive 20S proteasome (20Sc). Unlike 20Sc, 20Si hydrolyzes the carboxyl-terminus of PI31 and this effect contributes to the reduced inhibitory activity of PI31 towards 20Si. These results demonstrate unexpected differential interactions of PI31 with 20Sc and 20Si and document their functional consequences.

## INTRODUCTION

The 20S proteasome (also known as Core Particle) is an evolutionarily conserved protease complex consisting of four axially stacked heptameric rings (*1*). In eukaryotes, each of the two identical outer rings contains seven different α-type subunits (α1-7) and each of the two identical inner rings contains seven different β-type subunits (β1-7) (*2*). Three subunits of each inner ring (β1, β2, and β5) feature N-terminal threonine residues that function as catalytic nucleophiles. These subunits display different specificities for peptide bond hydrolysis (cleavage after acidic, basic/neutral, and small hydrophobic residues, respectively) (*3*). Cells specializing in immune function or cells exposed to certain cytokines, such as interferon-γ, selectively express genetically distinct forms of each catalytic subunit (β1i, β2i and β5i, respectively) that are selectively incorporated into otherwise compositionally identical 20S proteasomes termed “immuno-proteasomes” (*4*). The β1i and β5i subunits have different substrate specificities (cleavage after small hydrophobic and bulky hydrophobic residues, respectively) relative to their β1c and β5c counterparts because of differences in their substrate binding pockets (*5*). These catalytic features of 20Si favor production of antigenic peptides for presentation on class I major histocompatibility complexes (*5–7*). Immuno-proteasomes may also participate in cellular responses to oxidative and other types of stress (*8, 9*).

Regardless of isoform, the collective action of the 20S proteasomes’ multiple, catalytically diverse active sites has the potential to extensively degrade the varied complement of cellular proteins to amino acids and short peptides, a feature consistent with the proteasome’s role as the predominant system for intracellular protein degradation (*10, 11*). Several structural properties of 20S proteasomes, however, normally restrict this proteolytic function. First, the catalytic sites line the surface of a hollow chamber in the interior of the complex (*2*). This architecture, a consequence of the abutting β subunit rings, physically sequesters the catalytic sites from potential substrates. Access to the catalytic chamber requires substrates to pass through narrow 13 angstrom pores in the center of the outer α rings (*12*). These pores act as filters that restrict substrates to short peptides and unstructured polypeptide chains. Second, regardless of their size or structure, potential substrates can be blocked from pore transit by N-terminal extensions of multiple α-subunits that lay across and occlude the pore. Pore occlusion appears to be the constitutive state of cellular 20S proteasomes, thereby rendering them catalytically inert (*13*). Physiologic activation of 20S proteasomes occurs when specific regulatory proteins bind to their outer α rings (*14*). Higher eukaryotes contain multiple regulators including the 19S/PA700 regulatory complex, PA28αβ, PA28γ and PA200 and p97 (*15*). Despite their different physiologic roles and exact mechanisms of action, each of these regulators renders the catalytically inert 20S proteasome competent for substrate degradation by inducing conformational changes that result in retraction of the occluding α subunit peptides and opening of the substrate access pore (*14, 16–19*).

PI31 (<u>P</u>roteasome Inhibitor of <u>31</u>,000 Da) was identified as an *in vitro* protein inhibitor of the constitutive 20S proteasome (*20, 21*). Several recent high-resolution cryo-EM structures of 20S-PI31 complexes reveal the structural basis for this effect by showing that the intrinsically disordered proline rich carboxyl-terminus of PI31 enters the central degradation chamber of 20S and interacts directly with each of three different catalytic β subunits while the globular N-terminal FP domain remains on the exterior surface of the α-rings (*22–24*). Because one PI31 monomer can enter each end of the 20S cylinder, all six catalytic subunits are engaged and inhibited. Thus, unlike 20S activators that bind to the surface of the α-subunit rings and induce opening of the substrate access gate, PI31 passes through the pore of the α-subunit rings and directly affects function of the catalytic sites. The PI31-20S structures also suggest a mechanism by which PI31 itself escapes proteolysis (*22, 23*). The mammalian PI31-20S complex was assembled *in vitro* from purified PI31 and the constitutive isoform of latent 20S (*23*). This raises questions about how PI31 gains access to the catalytic sites and how it might interact with the structurally distinct β subunits of 20Si. In light of known differences between structural and catalytic features of 20Sc and 20Si proteasomes, we sought to compare the relative interactions of PI31 with these different proteasome isoforms. We report that 20Si is inhibited less effectively than 20Sc by PI31. Moreover, unlike 20Sc, 20Si extensively hydrolyzes PI31’s carboxyl terminus, a feature that may contribute to its lower degree of inhibition. These results demonstrate unexpected differential interactions of PI31 with 20Sc and 20Si and document their functional consequences.

## RESULTS

### Purified catalytically latent 20Sc and 20Si proteasomes are activated by gate-opening agents

Because PI31’s mechanism of inhibition requires passage of PI31 though the gate formed by 20S α subunit rings, we characterized the latent nature of 20S proteasomes used for subsequent experiments. We purified constitutive and immuno-forms of bovine 20S proteasome (20Sc and 20Si, respectively) to homogeneity, as documented by their migration as single bands upon native PAGE, and confirmed their identities by the differential composition of their unique catalytic β subunits (Figure 1). The catalytically latent nature of each enzyme was confirmed by the ability of several different agents to greatly stimulate their activities (Figures 1 and 2). Thus, latent 20Sc and 20Si featured low proteolytic activity in multiple orthogonal *in vitro* assays of proteasome function, including the hydrolysis of AMC-linked peptides (both in soluble assays and in zymography after native PAGE) and of intrinsically disordered proteins such as casein and α-synuclein (Figures 1 and Figure 2A and 2B). The catalytic subunits of latent 20Sc and 20Si were also labeled to low extents with the fluorescent proteasome activity probe, Me_4_BodipyFL-Ahx_3_Leu_3_-VS (Figure 2C). 20Sc and 20Si activities in these assays were increased (5-15 fold in various assays) by non-denaturing concentrations of SDS (*25*), the physiologic proteasome activator protein PA28 (*26, 27*), and carboxylbenzyl-tyrosine-alanine (z-YA), a dipeptide that mimics the HbYX activation motif of other physiologic proteasome activators such as the Rpt subunits of the 19S/PA700 regulatory particle (*28*). Each of these activating molecules promotes opening of the substrate access gate of latent proteasomes, thereby increasing substrate supply to the catalytic sites within the proteasome’s central catalytic chamber (*16, 28*). These results confirm the latent nature of the purified 20Sc and 20Si proteasomes and demonstrate their activation by mechanisms that involve gate opening.

**Figure 1.**
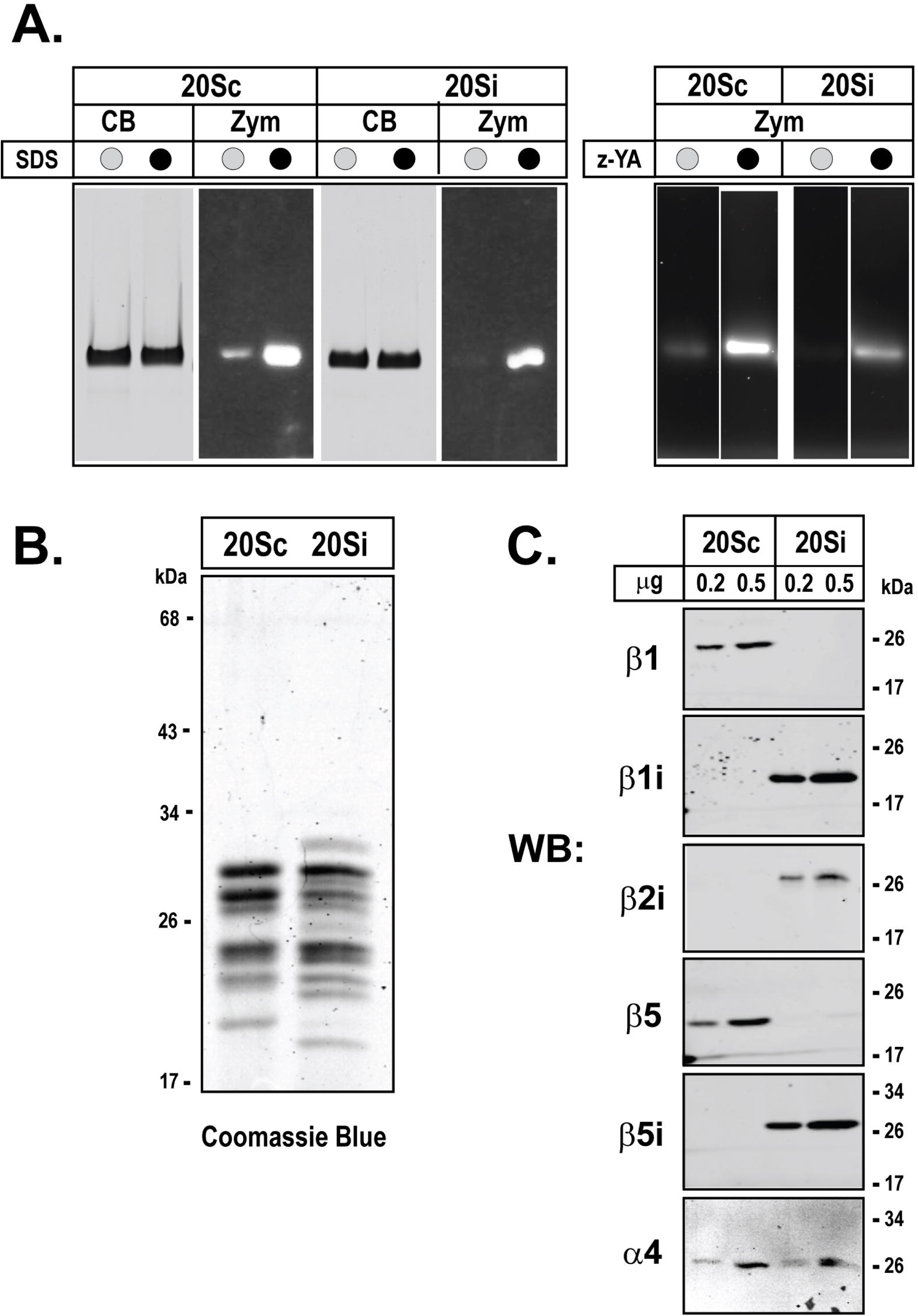
Characterization of purified bovine 20Sc and 20Si proteasomes. 20Sc and 20Si proteasomes were purified from bovine red blood cells and spleen, respectively as described under Experimental Procedures. *Panel A (left)*. Purified constitutive and immuno-20S proteasomes (5 μg/lane) were subjected to native PAGE on 3-8% acrylamide gels and either stained with Coomassie Blue-R250 (CB) or subjected to zymography (Zym) using Suc-LLVY-AMC substrate in the presence (•) or absence (O) of 0.03% SDS. *Panel A (right).* 20Sc and 20Si (5 μg/lane) were preincubated in the presence (•) or absence (O) of 5 mM z-YA and subjected to native PAGE and zymography using Suc-LLVY-AMC substrate. *Panel B*. Purified latent 20Sc and 20Si (5 μg/lane) were subjected to SDS-PAGE and stained with Coomassie Blue-R250. *Panel C.* Purified 20Sc and 20Si (0.2 μg and 0.5 μg) were subjected to western blotting (WB) with antibodies specific for indicated subunits.

**Figure 2.**
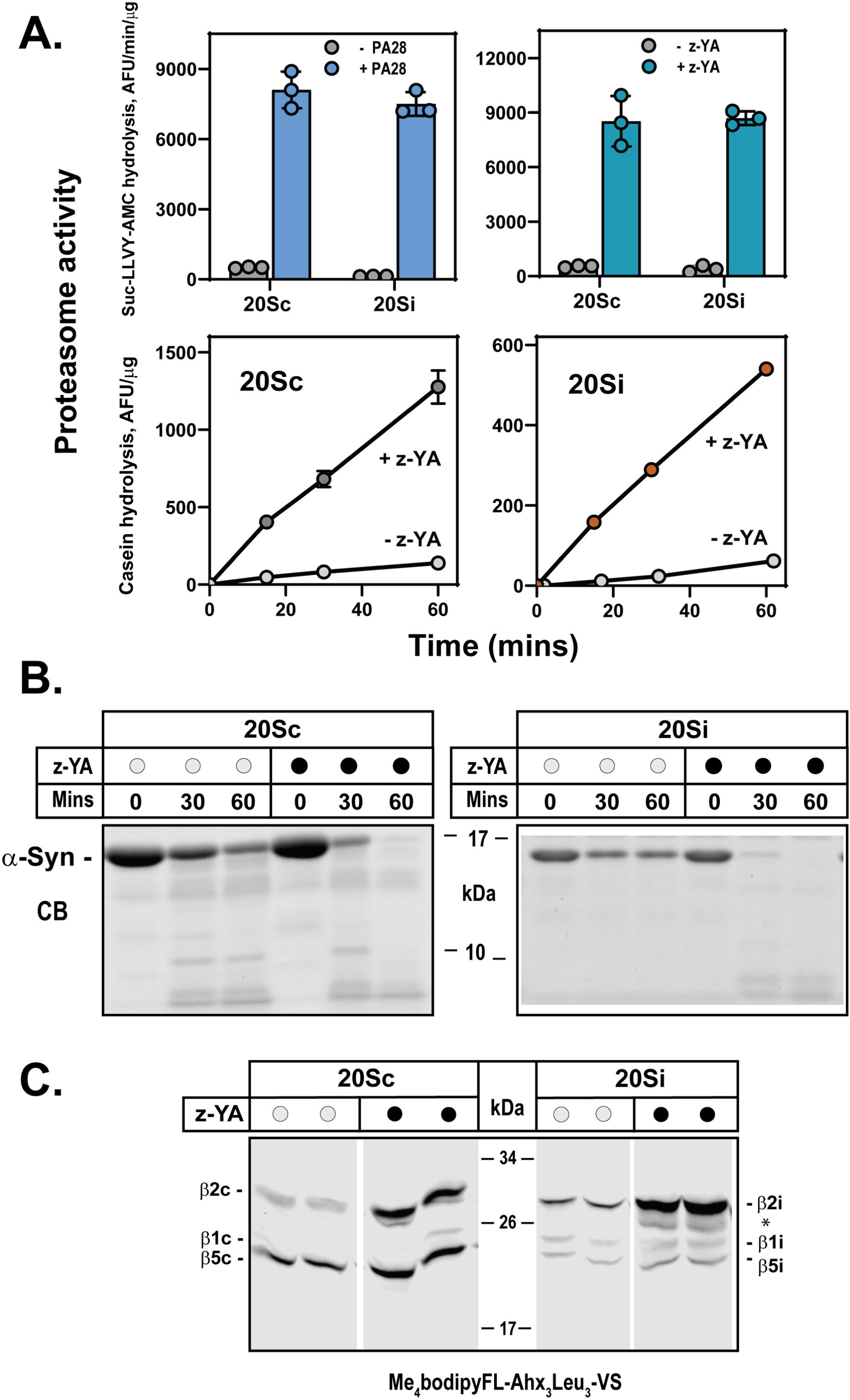
Purified 20Sc and 20Si are latent proteasomes. *Panels A and B*. Purified 20Sc and 20Si were assayed for proteasome activity with Suc-LLVY-AMC or casein substrates as described under Experimental Procedures in the presence or absence of PA28 or 5 mM z-YA. Data points represent mean values (± standard deviation) of triplicate assays. Similar results were obtained in at least three independent assays. *Panel B*. Purified 20Sc and 20Si (4.5 μg) were assayed for degradation of α-synuclein (α-Syn) in the presence (•) or absence (O) of 5 mM z-YA. At indicated times, reactions were terminated and samples were subjected to SDS-PAGE and stained with Coomassie Blue R-250 (CB). *Panel C.* Purified 20Sc and 20Si were preincubated in the presence (•) or absence (O) of 5 mM z-YA prior to treatment with 500 nM Me_4_BodipyFL-Ahx_3_Leu_3_-VS for 30 mins at 37°C. Samples were subjected to SDS-PAGE and gels were imaged as described under Material and Methods. Results for replicate incubations are shown. Similar results were obtained in three independent experiments. * indicates a band whose identity cannot be reliably established.

### PI31 inhibits immuno-20S proteasome less efficiently than it inhibits constitutive 20S proteasome

Our recent cryo-EM structure of mammalian PI31 in complex with 20Sc revealed specific binding interactions between residues of PI31 and β subunits within the catalytic chamber of the proteasome. These interactions include those between PI31 and the catalytic threonine residues that account for PI31 inhibition of the proteasome. Because 20Sc and 20Si differ solely by the identity of their catalytic β subunits, we sought to compare the PI31 inhibition of these proteasomes in the functional assays described above. As expected from previous work, PI31 inhibited 20Sc-catalyzed hydrolysis of both AMC-linked peptides and natively disordered proteins (Figure 3A and 3B). PI31 also inhibited the labeling of catalytic βc subunits by Me_4_BodipyFL-Ahx_3_Leu_3_-VS (*20, 21*) (Figure 3C). Although inhibition was observed against both latent and z-YA-activated 20Sc in each of these assays, the degree of inhibition against basal activity of the latent 20Sc was variable (20-80% inhibition) among repeated independent experiments. This variability was most prominent at low molar ratios of PI31:20S (1:1 – 5:1) and may reflect variability in the extent to which the C-terminus of PI31 is able to transit the substrate access pore of latent 20S to reach the catalytic sites. Additional variability likely results from calculations of percent inhibition based on two very low levels of proteasome activity. In contrast activated 20Sc was inhibited consistently and typically to a greater degree than latent 20Sc at given PI31 concentrations. PI31 also inhibited both latent and z-YA-activated 20Si in these same assays (Figures 3A, 3B and 3C). As with 20Sc, the degree of PI31 inhibition of basal substrate hydrolysis by latent 20Si was variable, but was reliably demonstrated with z-YA-activated 20Si, albeit to a lesser magnitude than for z-YA-activated 20Sc. PI31 also blocked labeling of each βi subunit of latent 20Si by Me_4_BodipyFL-Ahx_3_Leu_3_-VS (Figure 3C). In contrast, PI31 failed to block labeling of any βi subunit of z-YA-activated 20Si. Surprisingly, PI31 promoted a 4-fold increase in labeling of the β5i subunit and a lesser, but consistently increased labeling of the β1i subunit the form of 20Si (Figure 3C). The mechanism of this unexpected effect is unclear, but may represent an allosteric activation of β5i and β1i by PI31. These effects may contribute to the diminished PI31 inhibition of proteasome activity in substrate hydrolysis assays by activated 20Si (see Discussion). Nevertheless, PI31 inhibited 20Si to a lesser degree than 20Sc, regardless of proteasome activation state. Thus, the IC50 of PI31 for 20Si in both latent and activated states was over 10-fold greater than that for 20Sc (Figure 3D). Collectively, these results indicate that PI31 interacts differently with the unique catalytic β subunits of 20Sc and 20Si and that this difference results in attenuated PI31 inhibition of 20Si.

**Figure 3.**
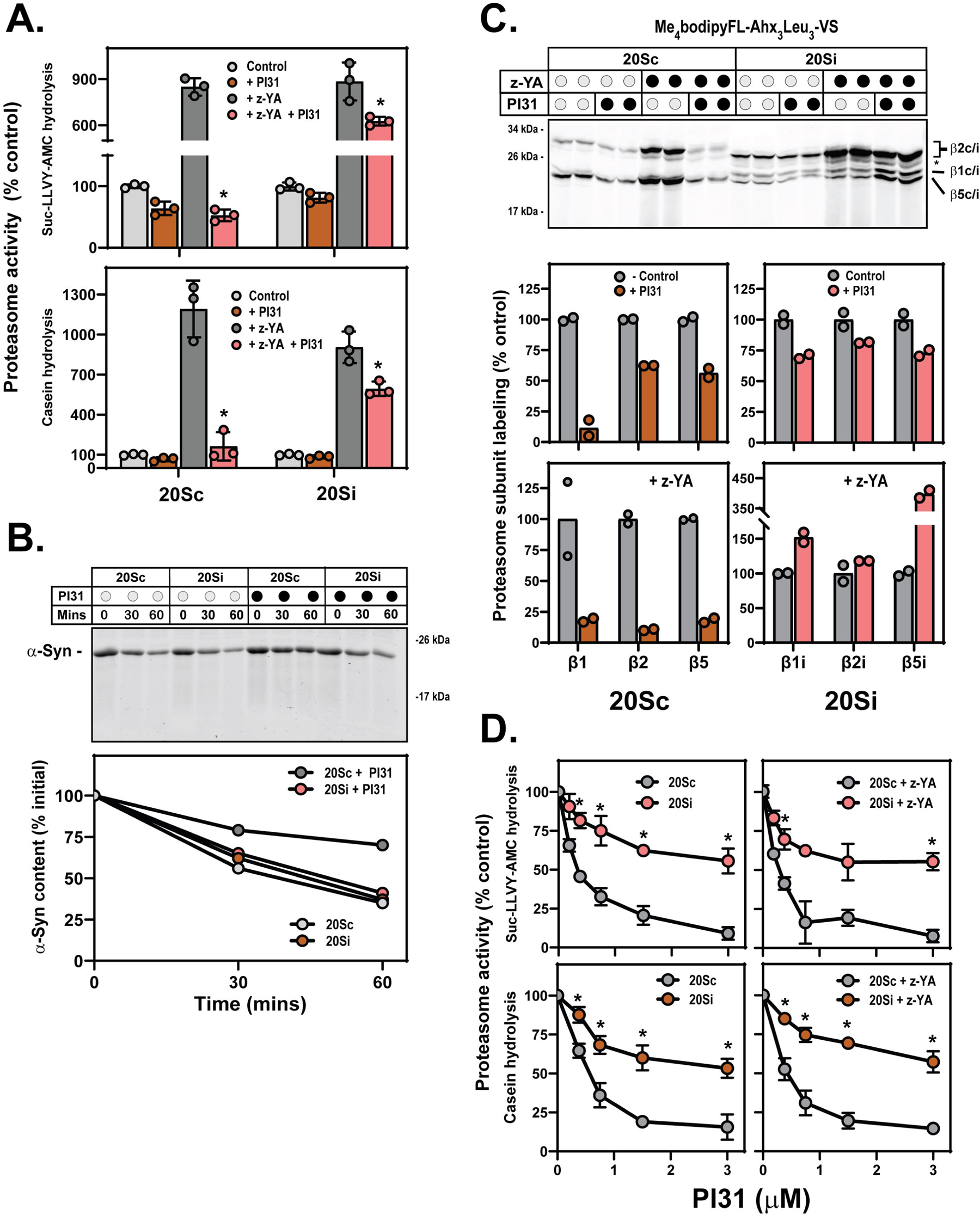
PI31 inhibition of 20Si is attenuated compared to that of 20Sc. PI31 inhibition of latent and z-YA-activated 20Sc and 20Si activities was compared in multiple assays of proteasome function. *Panel A*. *Effect of PI31 on latent and z-YA-activated hydrolysis of Suc-LLVY-AMC (upper) and casein (lower)*. Mean values for rates of substrate hydrolysis for triplicate assays were determined as described under Experimental Procedures. Values for activity in the absence of z-YA and PI31 were set to 100% (Control) and other values were calculated as a percentage of Control, ± standard deviation. Differences among results of different assay conditions were analyzed by 3-way ANOVA. Pair-wise differences were analyzed post-hoc by Tukey’s HSD. PI31 significantly inhibited z-YA activated 20Sc- and 20Si-catalyzed hydrolysis of both substrates (20Sc, p < 0001; 20Si, p = 0.0193). PI31 inhibition of 20Si was significantly attenuated compared to that of 20Sc for both substrates (p < 0.0001, denoted by *). *Panel B*. z-YA-activated 20Sc and 20Si hydrolysis of α-synuclein was determined in the presence (•) or absence (O) of PI31 (20S, 150 nM; PI31, 500 nM). At indicated times, samples were subjected to SDS-PAGE and stained with Coomassie Blue R-250. α-synuclein content was quantified using Image Studio software (LiCOR). Values at time 0 were set to 100% and other values within each group are represented as a percentage. Similar results were obtain in five independent experiments. *Panel C*. Me_4_BodipyFL-Ahx_3_Leu_3_-VS labeling of 20Sc and 20Si in the presence (•) or absence (O) of 5 mM z-YA and/or PI31. Labeling was conducted as described in Experimental Procedures and quantified using Image Studio software (LiCOR). Mean values for replicates of labeling of each subunit in the absence of z-YA and PI31 were set to 100%. Other values within that group are represented as percentages of that value. Similar results were obtain in two other independent experiments. *Panel D*. Purified 20Sc and 20Si were assayed for hydrolysis of Suc-LLVY-AMC and casein in the presence or absence of 5 mM z-YA at indicated concentrations of PI31. Mean values for rates of hydrolysis of triplicate assays in the absence of PI31 were set to 100% and other values were determined as a percentage of that value. Differences in activity between 20Sc and 20Si were analyzed by repeated measures of 2-way ANOVA. Pair-wise, concentration-matched differences in activity between 20Sc and 20Si were analyzed by Šídák’s multiple comparisons test, and significant differences are indicated at p ≤ 0.05 by (*) for respective PI31 concentrations. Similar results were obtained in four independent experiments.

### 20Si proteasome hydrolyzes PI31

The high-resolution structure of the mammalian PI31-20Sc complex (*23*) and a similar structure of the complex from yeast (*22*) reveal both a mechanism for proteasome inhibition via interaction with catalytic β subunits and an explanation for how PI31itself escapes proteolysis. The different degrees of 20Sc and 20Si inhibition by PI31 described above might be a manifestation of differences in either or both of these features. Therefore, we compared the susceptibility of PI31 to 20Sc and 20Si-catalyzed proteolysis by monitoring the fate of intact PI31 during incubation with each proteasome subtype in the presence and absence of z-YA, but in the absence of an exogenous proteasome substrate. In accord with our original findings, PI31 was largely resistant to hydrolysis by both latent and activated 20Sc (*20*) (Figure 4). A minor loss of intact PI31was sometimes detected after extended incubations, but even this minimal degree of degradation was not observed consistently. Intact PI31 content was likewise lost to a minimal extent when incubated with latent 20Si. As with the minimal effect of PI31 on latent 20Si activity, this result may reflect restricted transit of PI31 through the predominately closed substrate access pores. In contrast, PI31 was extensively hydrolyzed by z-YA-activated 20Si (Figures 4A and 4B). The loss of intact PI31 was completely blocked by MG132, confirming that the reduced PI31 content was caused by proteasome activity (Figure 4A, *right panel*).

**Figure 4.**
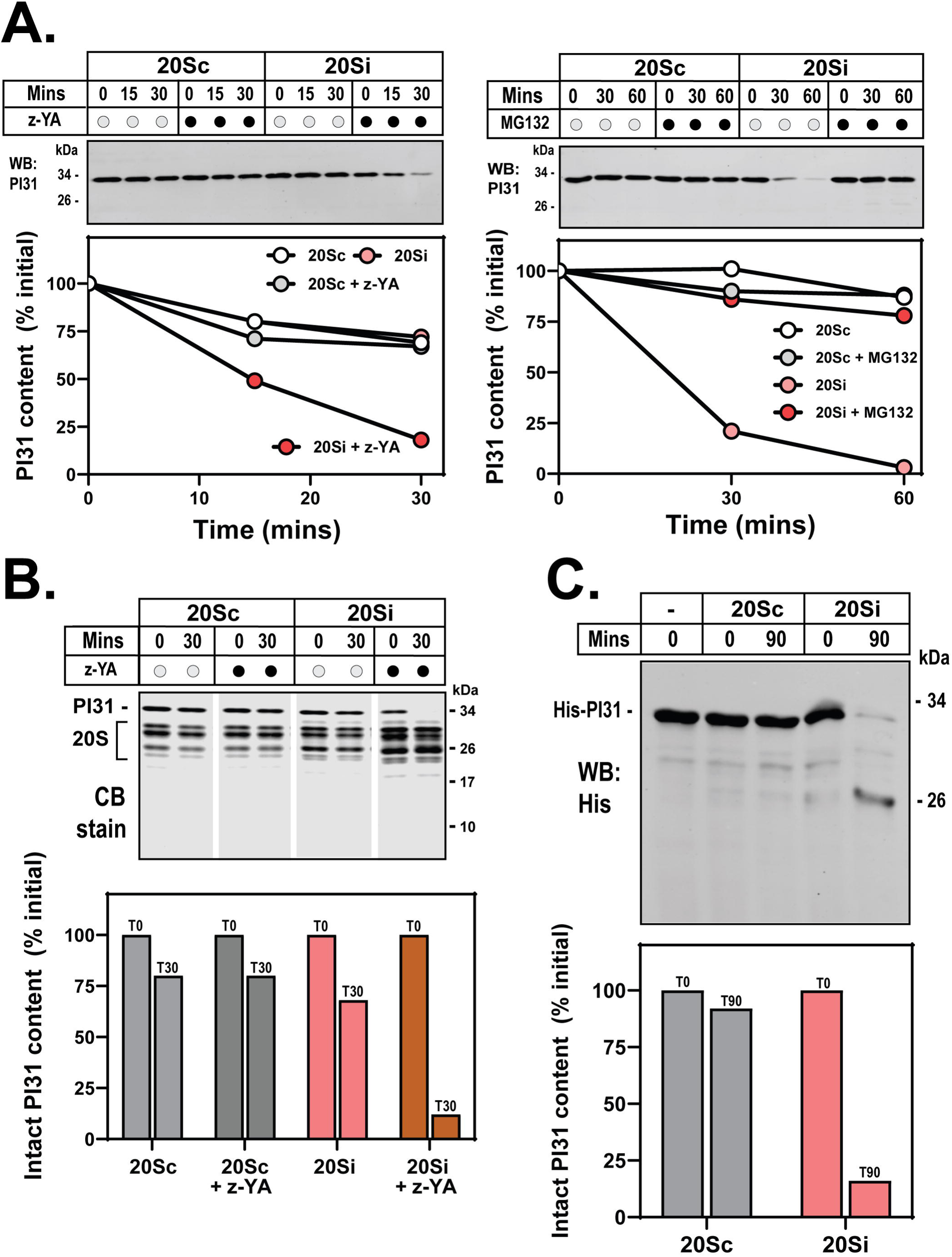
20Si selectively hydrolyzes PI31. *Panel A, (left)*. Purified latent (O) and z-YA-activated (•) 20Sc and 20Si (200 nM) were incubated with PI31 (1 μM) at 37° C for indicated times. Reactions were terminated with SDS samples buffer. Samples were subjected to SDS-PAGE and western blotting with an antibody against the C-terminus of PI31. PI31 bands were quantitated with Image Studio software (LiCOR). Values within each group are expressed as a percentage of the initial value. Similar results were obtained in five independent experiments. *Panel A (right)*. z-YA-activated 20Sc and 20Si were incubated as described above in the presence (•) or absence (O) of 0.5 mM MG132 prior to incubation with PI31, as in Panel A, right. *Pane B (upper)*. As in Panel A, but the gel was stained with Coomassie Blue R-250 (CB). *Panel C*. As in Panel A, but gel was western blotted with anti-His antibody. *Panel B (lower).* Quantification of PI31 bands from upper panel. Similar results were obtained in three independent experiments. *Panel C (upper).* PI31 was incubated with z-YA-activated 20Sc or 20Si as in Panel A. After 90 mins reactions were terminated with SDS samples buffer and subjected to western blotting using an anti-His antibody. *Panel C (lower)*. Quantification of intact (T0) PI31 content from upper panel. Similar results were obtained in four independent experiments.

The topology of PI31 hydrolysis by z-YA-activated 20Si was assessed using several different monitors of PI31 content, including western blotting with antibodies selective for either the N- or C-terminus of the protein (Figures 4A and 4C). z-YA-activated 20Si promoted loss of the C-terminus without accumulation of detectable fragments . In contrast, a PI31 degradation product of about 24 kDa containing the N-terminus accumulated during the same incubation conditions (Figure 4C). These results are consistent with the known orientation of PI31 within the PI31-20S structure and suggest that the last ∼70 amino acids (approximately 8000 Da) of the C-terminus of PI31 are cleaved from the protein by activated 20Si. We hypothesize that the folded N-terminal domain of PI31 is unable to pass through the z-YA-opened pore, thereby limiting the extent to which the remaining ∼50 residues (i.e., residues ∼ 152-200) of PI31’s C-terminal domain can be hydrolyzed further. To test this hypothesis, we assessed 20S-catalytzed hydrolysis of a series of PI31 mutants with progressively greater C-terminal truncations (Figure 5A). We reasoned that C-terminal PI31 sequences of insufficient length to reach the proteasome’s catalytic sites should be unavailable for proteolysis. Accordingly, PI31^1-181^ and PI31^1-193^, neither of which had inhibitory activity, were refractory to hydrolysis by both 20Sc and 20Si (Figures 5B and 5C). In contrast, PI31^1-244^, which extends into the catalytic chamber and exerts a modest inhibitory effect on 20Sc, was hydrolyzed by activated 20Si but only to a minor degree by 20Sc (Figure 5C). This hydrolysis resulted in the accumulation of a 24 kDa fragment that was indistinguishable from that produced by hydrolysis of full-length PI31 (PI31^1-271^) (Figure 5B). Collectively, these results show that PI31 is selectively hydrolyzed by 20Si and indicate that hydrolysis of wild-type PI31 is limited to those resides that extend to the catalytic chamber.

**Figure 5.**
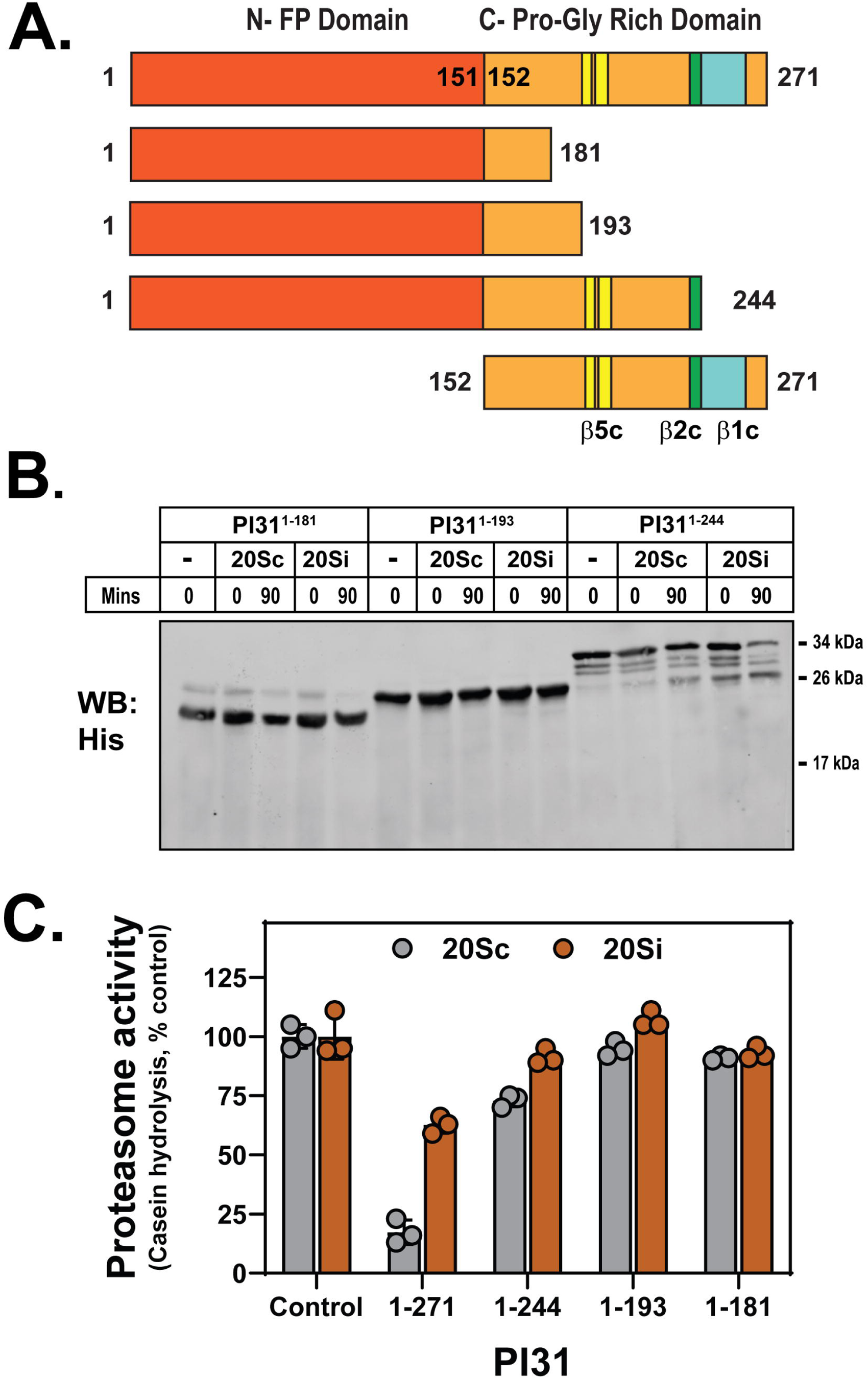
Topology of PI31 susceptible to hydrolysis by 20Si. *Panel A*. Schematic representation of domain structure of wild type (1–271) PI31. The positions of regions of PI31shown previously to interact with the β5 (yellow), β2 (green), and β1 (blue) catalytic residues of the proteasome are indicated. Structures of PI31 mutants with C-terminal truncations are shown for comparison. *Panel B.* Indicated mutants of PI31 were expressed as N-terminal His-tagged recombinant proteins in *E. coli* and purified as described under Experimental Procedures. Each mutant PI31 was incubated at 37°C for indicated times with z-YA-activated 20Sc or 20Si. Samples were subjected to western blotting with anti-His antibody. Similar results were obtained in two independent experiments. *Panel C*. Wild-type PI31 and indicated PI31 mutants (1.5 μM) were assayed for inhibition of proteasome activity of casein hydrolysis by 20Sc and 20Si (150 nM), as described under Experimental Procedures. Mean values for triplicates assays of proteasome activity of each proteasome were set at 100% and other values were expressed as a corresponding percentage. Differences in relative proteasome activity for 20Sc and 20Si incubated with different truncations of PI31 were analyzed by 2-way ANOVA. 20Sc was significantly inhibited by PI31^1-271^ (Tukey’s HSD; adj. p < 0.0001) and PI31^1-244^ (Tukey’s HSD; adj. p < 0.0001). 20Si was significantly inhibited by PI31^1-271^ (Tukey’s HSD; adj. p < 0.0001). PI31^1-271^ inhibition of 20Si was significantly less than that of 20Sc (Tukey’s HSD; adj. p < 0.0001). Similar results were obtained in two independent experiments.

The structure of the PI31-20Sc complex identified specific interactions between residues of the C-terminus of PI31 and individual βc subunits, including their catalytic threonines (*23*). The most N-terminally-positioned PI31 residue to make such a contact was aspartic acid 197, which forms a hydrogen bond with the catalytic threonine of β5c. 20Si cleavage of PI31 at (or near) this site would produce an N-terminal PI31 fragment predicted to be approximately 23,500 Da, a value in excellent accord with that of the experimentally detected fragment. To gain additional detail about the cleavage site responsible for production of the N-terminal PI31 fragment, we purified the 20Si-generated fragment and subjected it to mass analysis by intact protein mass spectrometry (Figure 6). The intact mass of full-length HisPI31 (not incubated with 20Si) was 31,804.22 Da (Figure 6B), a value that differs from the calculated mass of HisPI31 (31,936.02 Da) by 131.80 Da and is likely explained by the loss of the N-terminal methionine associated with co-translational methionine aminopeptidase activity during recombinant protein expression. Intact mass analysis of the purified HisPI31 20Si-generated degradation product had a mass of 23,693.87 Da (Figure 6D). This value differs from that of full-length HisPI31 by 8,110.35 Da, a value that corresponds to a loss of 78 residues from the C-terminus and identifies the cleavage site between V193 and G194 (see Discussion). This assignment is also consistent with the relative susceptibilities of various PI31 truncation mutants to hydrolysis shown in Figure 5B.

**Figure 6.**
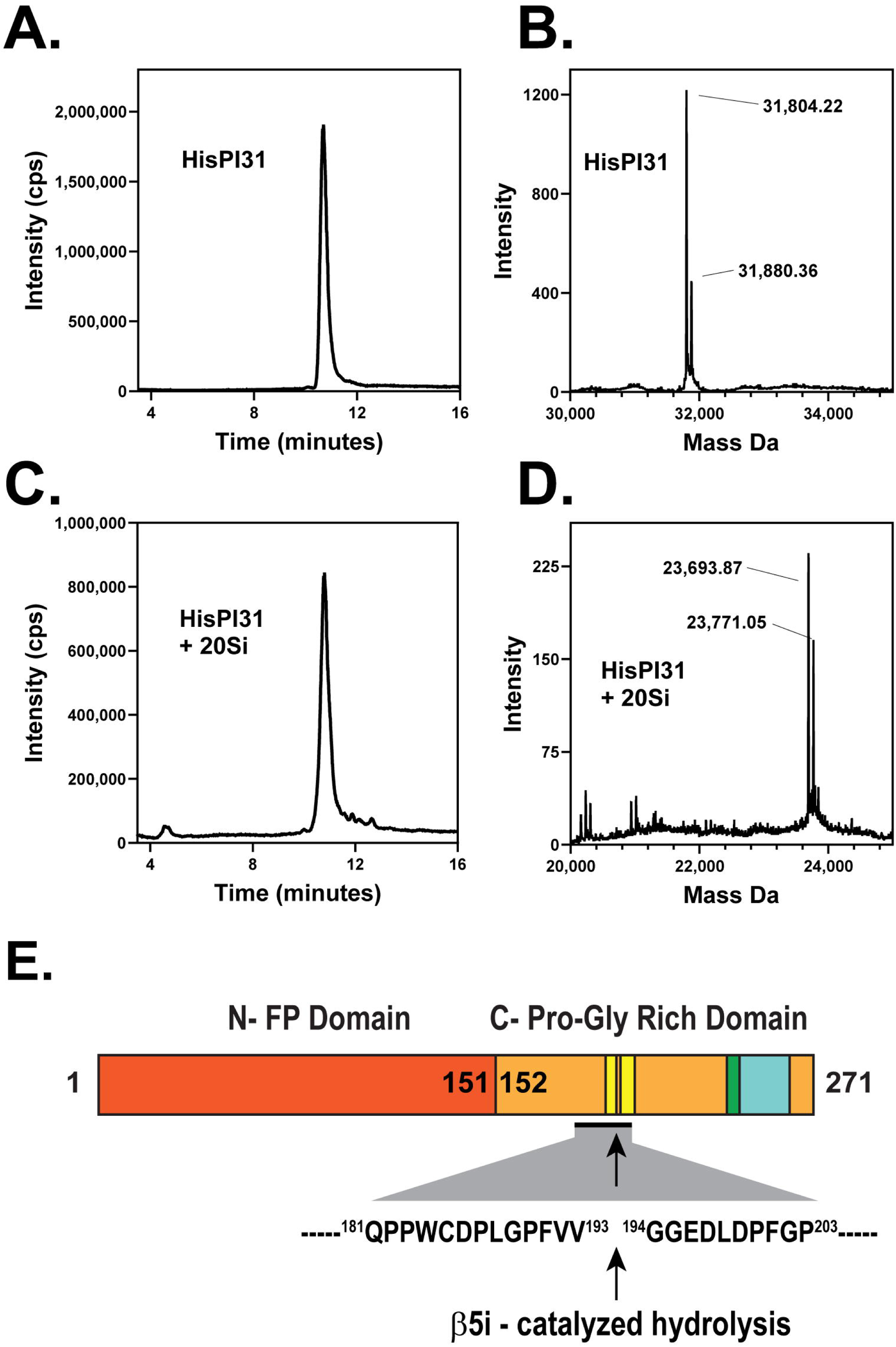
Identification of 20Si-catalyzed cleavage site of PI31 by intact mass analysis. Intact mass spectrometry was performed on recombinant His-PI31 before and after incubation with z-YA-activated 20Si. *Panel A.* Reverse phase chromatogram of purified, full-length HisPI31 injected for LC-MS. *Panel B.* Mass spectra of full-length HisPI31 from Panel A. *Panel C.* Reverse phase chromatogram of purified, HisPI31 20Si degradation product injected for LC-MS. *Panel D.* Mass spectra of HisPI31 20Si degradation product from Panel C. *Panel E*. Representation of domain structure of PI31, and the 20Si hydrolysis site derived from comparisons of Panels B and D. The dominant mass peaks before and after incubation with 20Si (*Panel B vs. D*) differ by 8,110.35 Da, a value consistent with the C-terminal 78 residues of PI31.

The folded N-terminal FP domain of PI31 (residues 1-151) appears to spare proteolysis of those PI31 residues of the C-terminal domain that span the distance between the open gate and the centrally located catalytic sites (residues from ∼S152 – V193). To further test this assertion, we incubated z-YA-activated 20Si with a PI31 mutant consisting of only the C-terminal proline-rich unstructured domain (His-PI31^152-271^). We confirmed previous results showing that this mutant was sufficient to inhibit activity of both the latent and activated forms of 20Sc (Figure 7A). In contrast PI31^152-271^ had appreciably less inhibitory activity against both latent and activated 20Si (Figures 7A and 7B). Analysis of PI31^152-271^ content by western blotting and Coomassie blue staining showed that the protein was selectively hydrolyzed by 20Si. The loss of the most N-terminal His-tagged portion of the peptide suggests that the hydrolysis of this peptide includes cleavage of portions (S152-V193) that were otherwise protected when linked with the N-terminal domain (Figure 7C). The partial inhibitory effect of the peptide on z-YA-activated 20Si may indicate that cleaved PI31 peptides exert inhibition while retained in the degradation chamber. Alternatively, inhibition of substrate hydrolysis may result from competition with PI31 hydrolysis. In sum, these results show that PI31 interacts differently with 20Sc and 20Si proteasomes. These differences appear to render the portion of PI31 within the catalytic chamber susceptible to hydrolysis by 20Si and this effect may contribute to attenuated inhibition by this form of the proteasome.

**Figure 7.**
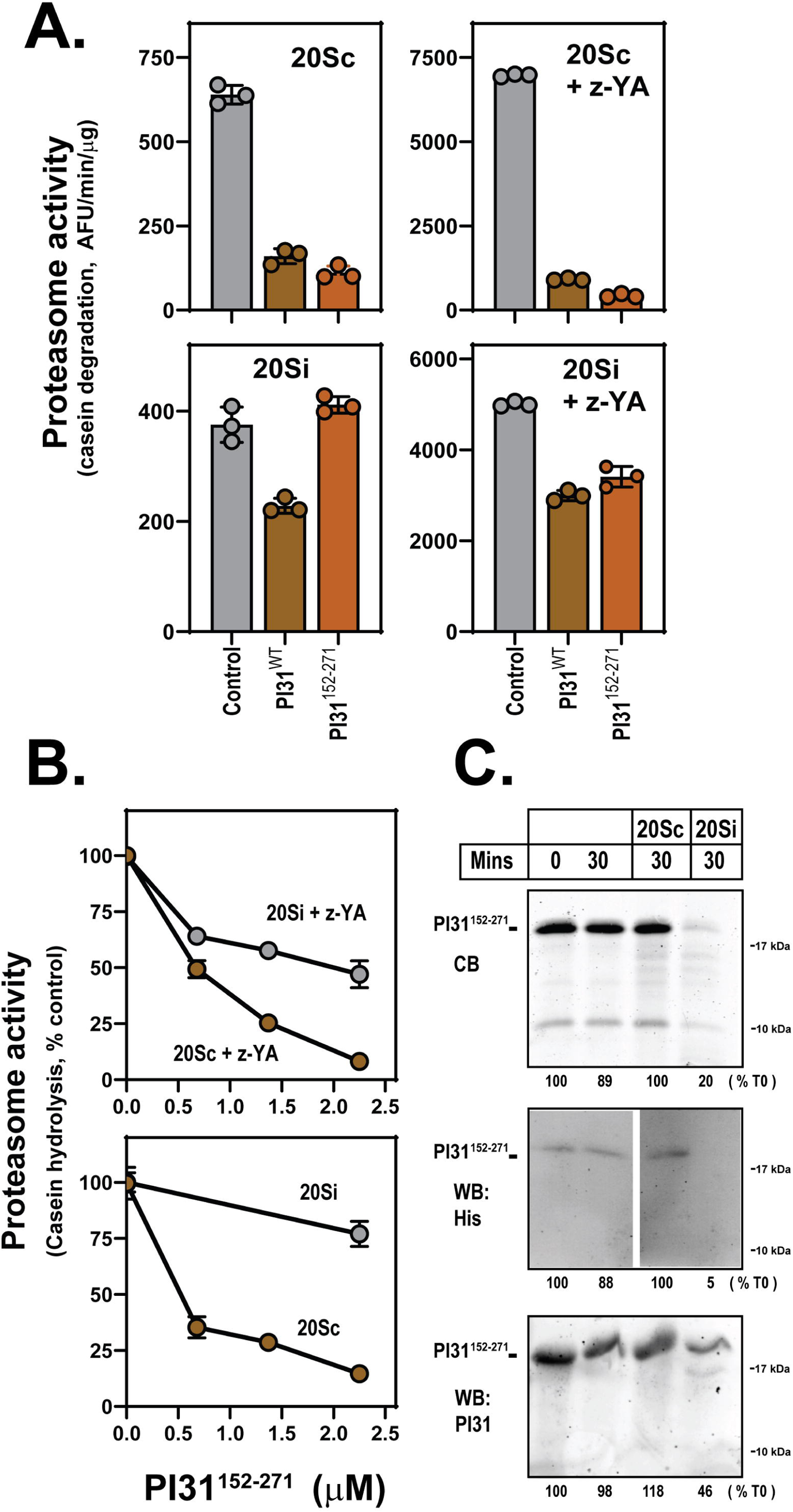
20Si selectively hydrolyzes the C-terminal domain of PI31. *Panel A.* Latent and z-YA-activated 20Sc and 20Si (150 nM) were assayed for proteasome activity in the presence or absence of full length PI31 (His-PI31^1-271^) and a mutant consisting of only the PI31 C-terminal domain (His-PI31^152-271^) (2.3 μM). Data are reported as mean values (± standard deviation) of triplicate assays for rates of casein hydrolysis. Similar results were obtained in three independent experiments. *Panel B.* Proteasome assays were conducted as in Panel A at indicated concentrations of His-PI31^152-271^. Proteasome activity in the absence of PI31 was set at 100% and other values are expressed as a percentage of that value. Similar results were obtained in three independent experiments. Panel C. 20Sc and 20Si (150 nM) were incubated with His-PI31^152-271^ (1.5 μM) for 30 mins at 37° C. Reactions were terminated with SDS sample buffer and subjected to SDS-PAGE for staining with Coomassie Blue R-250 (CB) or western blotting with antibodies against His (for the N-terminus) or for the C-terminal portion of the peptide. Similar results were obtained in three independent experiments.

### Hydrolyzed PI31 remains bound to 20S proteasome

To gain additional insight to the relative interactions of PI31 with 20Sc and 20Si proteasomes, we monitored formation of PI31-20S proteasome complexes by glycerol density gradient centrifugation. We preincubated wild-type and mutant PI31 proteins with either latent or z-YA-activated 20Sc or 20Si and analyzed the distribution of PI31 after centrifugation. In the absence of 20S, PI31 was detected exclusively in fractions containing slowly sedimenting proteins with low native molecular weights (Figure 8). After incubation with latent 20Sc, a portion of wild-type PI31 was identified in gradient fractions containing the proteasome, an indication complex formation. Activation of 20Sc with z-YA increased the binding of wild-type PI31, an effect that likely reflects increased passage of PI31 through the open pores of the activated proteasome. Most of the 20Sc-associated PI31 retained its native molecular weight, as determined by SDS-PAGE. In some experiments, such as shown in Figure 8, a small portion of the bound PI31 was detected as a PI31 hydrolysis product. Because this method cannot distinguish between proteasomes bound to one or two molecules of PI31, some of this hydrolysis may result from the action of uninhibited catalytic sites. Nevertheless, these findings confirmed that PI31 formed a largely hydrolysis-resistant complex with 20Sc. In contrast, no detectable complex was formed between latent 20Si and wild-type PI31. As with z-YA-activated 20Sc, most nearly all PI31 bound to z-YA-activated 20Si. However, this bound PI31 was extensively hydrolyzed and the major fragment was indistinguishable from the 24 kDa hydrolysis product detected in experiments described above. Preincubation of 20Si with the small molecule immunoproteasome inhibitor, ONX-914, prior to PI31-20Si complex formation resulted in a complex featuring completely intact PI31. Likewise, PI31 bound efficiently to 20Sc that had been treated with expoxomicin.

**Figure 8.**
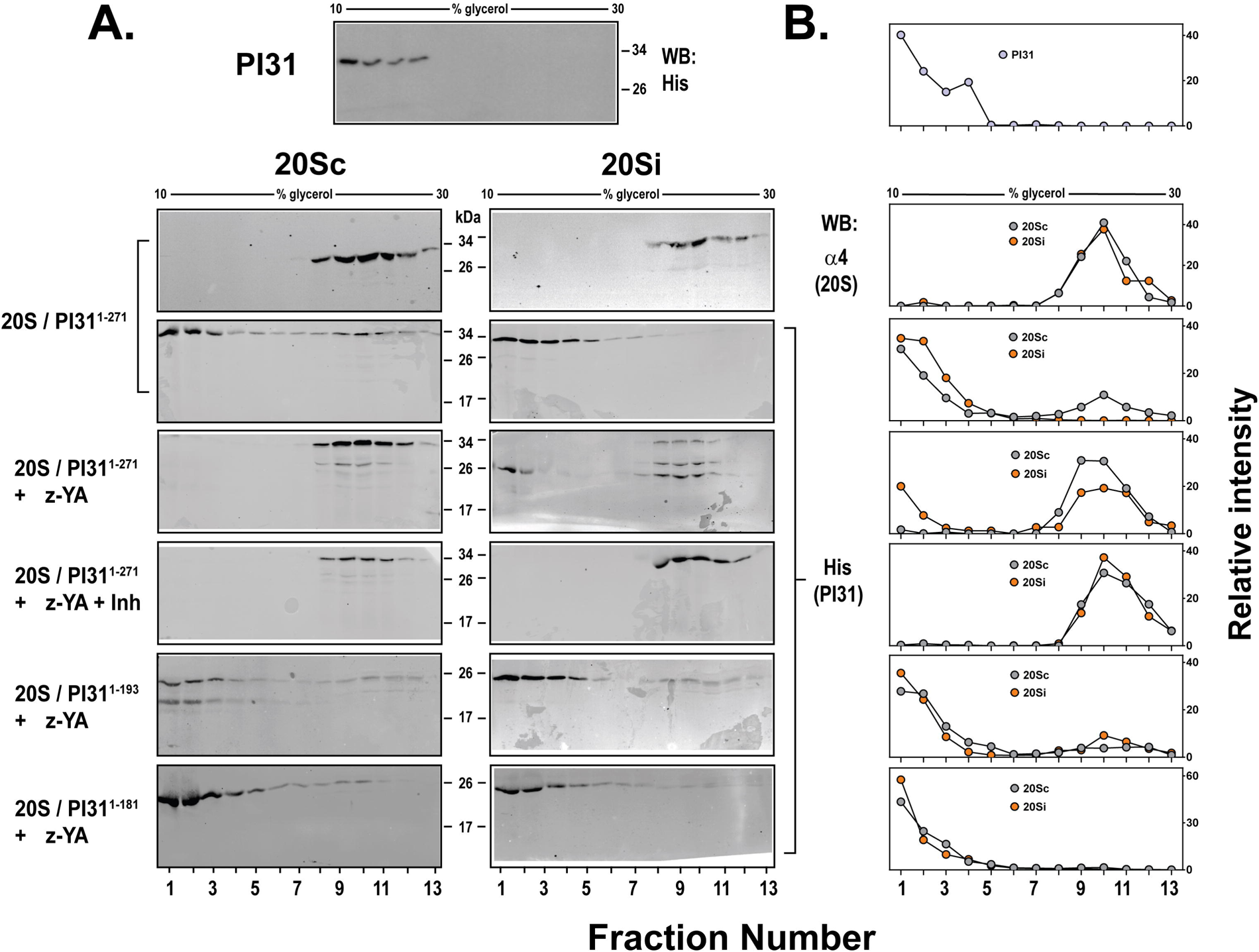
Structural determinants of PI31 for complex formation with 20Sc and 20Si. 20Sc or 20Si were preincubated with the indicated PI31 proteins and subjected to glycerol density gradient centrifugation as described under Experimental Procedures. *Panel A.* Distribution of His-PI31 was determined by western blotting of using anti-His tag antibody. Control gradients were performed for proteasome in the absence of PI31 using an antibody against the α4 subunit common to both 20Sc and 20Si and against PI31 in the absence of 20S. *Panel B*. . PI31 bands in Panel A were quantified in ImageStudio (LiCOR) for each fraction and expressed as a percentage of the sum total intensity for all fractions. In lanes where multiple bands appear, the area containing all bands was used for quantification Similar results were obtained in three independent experiments.

**Figure 9.**
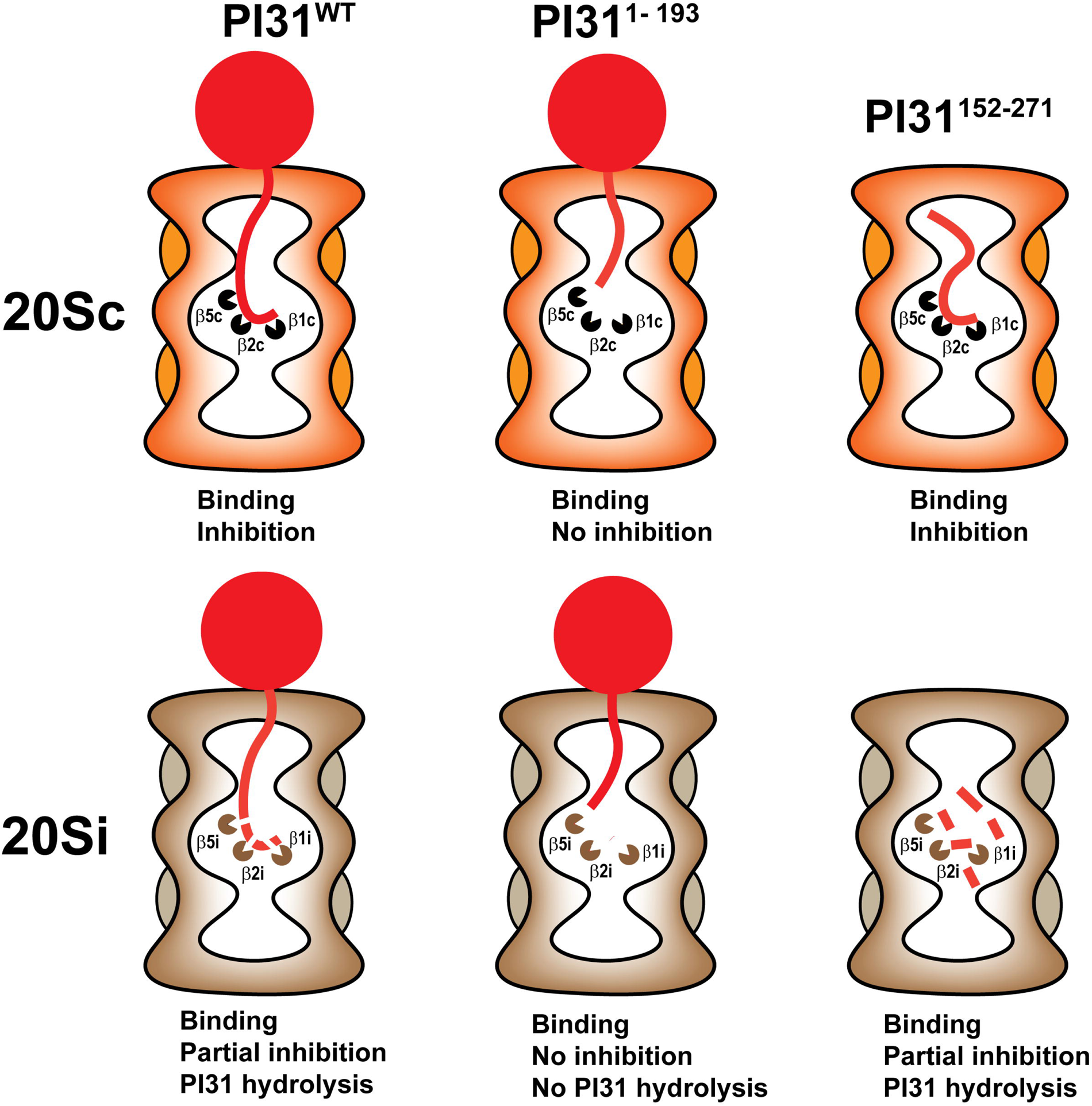
Model of PI31 binding to and hydrolysis by 20Sc and 20Si proteasomes. The intrinsically disordered C-terminal domain of PI31 enters the central chamber of 20Sc and 20Si proteasomes. PI31^1-271^ binds to βc catalytic sites to inhibit their function while escaping hydrolysis itself. Truncated PI31s with C-termini that cannot reach the catalytic sites PI31^1-<^ ^193^ feature no 20S inhibition and have reduced binding. PI31^1-271^ also enters the central chamber of 20Si but interacts with βi catalytic subunits in a manner that leads to reduced proteasome inhibition and PI31 hydrolysis. For clarity, the images depict a single molecule of PI31 interacting with three of the six catalytic subunits of 20S proteasomes. However, two molecules of PI31 can enter the proteasome from opposite ends of the cylinder and interact with all six catalytic subunits.

To further analyze determinants of PI31 binding to 20Sc and 20Si, we repeated the experiments described above using the C-terminally truncated PI31^1-181^ and PI31^1-193^ proteins (Figure 8). Although both mutants had detectable association with z-YA activated-20Sc and 20Si, in each case much less complex formation was obtained than with wild-type PI31 (Figure 8B). Based on results described above, neither of these 20S-associated proteins is expected to interact with the catalytic sites and accordingly neither was detectably hydrolyzed. These results suggest that PI31 residues of the C-terminal domain directly adjacent to the N-terminal FP domain (i.e. residues 152-193 that probably represent densities identified in the lateral antechambers of the 20S proteasome (*23*), contribute only minimally to the formation of stable PI31-20S complexes.

## DISCUSSION

Recently reported cryo-EM structures of PI31-20S complexes demonstrate that PI31 inhibits proteasome activity by binding directly to each catalytic threonine residue of 20Sc (*22–24*). PI31 binding occurs in a manner that also indicates a structural basis for resistance of PI31 to its own proteolysis, a feature confirmed here by direct tests. The current work demonstrates that PI31 inhibition of 20Si is significantly attenuated in relation to that of 20Sc and that PI31 undergoes 20Si-catalyzed degradation. Because 20Sc and 20Si have identical α subunits and are activated similarly by various gate-opening agents, it seems unlikely that they differ in their ability to allow passage of PI31 through their pores (*29*). Instead, these differences in PI31-induced inhibition and PI31 stability most likely result from the differences in the manner by which PI31 interacts with the unique catalytic β subunits of these proteasomes. Previous work has documented structural distinctions between the substrate binding pockets of β1c and β1i and between the substrate binding pockets of β5c and β5i. These distinctions account for the different substrate specificities of 20Sc and 20Si, as well as their preferential interactions with certain small molecule inhibitors (*29*). Accordingly, they may also account for the distinctions reported here for both PI31-mediated inhibition and PI31 stability. For example, the structure of the PI31-20Sc complex shows that aspartic acid residue 197 of PI31 forms a hydrogen bond with catalytic threonine of β5c at the S1’ position to inhibit its function while PI31 residues 194-212 are arranged in the substrate pocket in the reverse direction of that required for normal substrate hydrolysis (*23*). These specific arrangements may be lost or altered in the in larger pocket of the PI31-20Si complex, rendering PI31 less effective as a direct inhibitor and more susceptible to hydrolysis. Our results identify a site of PI31 cleavage between valine 193 and glycine 194. Notably, valine is a branched-chain, hydrophobic residue known to be preferentially hydrolyzed by β5i (*5, 30*).

PI31 inhibition was observed against basal activity of catalytically latent 20S proteasomes. However, degree of PI31 inhibition of basal activity of latent proteasome was highly variable among independent experiments when conducted at lower PI31 concentrations and required supra-stoichiometric levels of PI31 relative to 20S to observe reliably. These findings suggest that, like peptide and protein substrates, PI31 entrance into the catalytic chamber of latent 20S is highly restricted by the occluded α ring pores. In fact, formation of the 20Sc-PI31 complex reported for our cryo-EM structure required extended preincubation of the component proteins with 5-fold molar excess of PI31; even then, only an estimated 34% of proteasomes on the cryo-EM grids were classified with PI31 mass, a finding consistent with the inefficiency of complex formation with latent 20S (*23*). Thus, inefficient PI31-20S complex formation likely reflects the low probability of the open gate state of the latent proteasome, and PI31 may gain entrance to the proteasome only during transient gate openings (*31, 32*). In contrast, PI31 inhibited open-gated proteasomes more efficiently across a range of PI31 concentrations. Although gate opening is obligatory for entry of PI31 into the interior of the proteasome, PI31 must also transit through the antechamber to reach the catalytic sites. Although the structure of the PI31-20S complex revealed PI31 density in the 20S antechambers, these interactions appear to contribute only minimally to the stability of the complex.

The structurally dimeric nature of the 20S proteasome raises additional complications for interpreting kinetic features of PI31 inhibition. For example, reported 20S-P31 structures feature two molecules of PI31 bound to the proteasome, one entering from each end of the 20S cylinder (*22, 23*). In this state, all six catalytic sites are engaged and inhibited. However, proteasomes bound to a single molecule of PI31 would have three unengaged catalytic subunits. Although we assume that these subunits to be catalytically active, we know of no biochemical evidence that validates this assumption. Therefore, the unengaged catalytic sites might contribute to PI31 hydrolysis observed with 20Si and to a minor degree with 20Sc. Partial engagement of the catalytic subunits by PI31 might promote other effects on the non-engaged subunits with respect to substrate hydrolysis. Our surprising finding of increased activity-probe labeling of the β5i and β1i subunits in the presence of PI31, is consistent with this speculation. Finally, PI31 hydrolysis by 20Si may contribute indirectly to measures of PI31 inhibition of substrate hydrolysis via a competitive mechanism. In fact, we detect significant 20Si-catalyzed PI31 degradation during the hydrolysis of protein substrates such as casein and α-synuclein (unpublished data). These various issues highlight the complications of providing precise quantitative kinetic characterization of PI31 inhibition.

20S-catalyzed hydrolysis of PI31 was limited to PI31’s C-terminus and is consistent with the orientation of the molecule within the catalytic chamber. Our 20S-PI31 structure showed that the most N-terminal portion of PI31 in contract with a catalytic threonine was aspartic acid 197 (*23*). Cleavage of PI31 by β5i near this residue, as described above, would produce a fragment of about 23,000 Da. The experimentally determined mass of 23,693.87 Da for the cleavage product supports this interpretation. We hypothesize that further cleavage of this fragment is restricted because the N-terminal FP domain prevents transit of residual the C-terminal polypeptide to the catalytic sites. Our results showing the resistance of PI31 C-terminal mutants consisting of residues 1-193 and 1-183, but the lability of a mutant consisting of residues 152-271 to hydrolysis support such a hypothesis. In contrast, the PI31 polypeptide chain C-terminal to residue 193 (i.e., 194-271) appears to be extensively degraded, possibly via its interactions with the β2 and β1 catalytic subunits.

Although the physiologic significance of the *in vitro* effects reported here is unknown, these results suggest different cellular roles for PI31 against 20Sc and 20Si proteasomes. We note that previous work identified a striking difference of the cellular association of PI31 with 20Sc and 20Si proteasomes and concluded that PI31 formed preferentially formed complexes with 20Sc compared to 20Si (*33*). It is unclear whether this difference might be a consequence of the loss of 20Si-PI31 complexes caused by PI31 degradation. PI31 lability through a process of degradation may also provide a mechanism to reverse its inhibitory activity. Such a reversal has been reported for PI31-inhibited 20S proteasomes that transition to active 26S-like proteasomes after germination of dormant *Microsporidia* spores (*24*); a possible causal relationship between this transition and PI31 degradation has not been established. Other work has shown that overexpression of PI31 inhibited the cellular assembly of 20Si proteasomes in response to interferon-γ (*34*). Although these findings suggest a functional link between 20Si and PI31, a mechanism for this effect was not defined, and there is no obvious relationship between this phenomenon and the results described in the current work.

The results presented here focus on distinctions of PI31’s *in vitro* action as an inhibitor of 20Sc and 20Si. However, as noted previously, there is little direct evidence for the physiologic significance of this function (*35*). Canonical views of cellular proteasome function involve activation and regulation of latent 20S proteasomes by activator complexes such as 19S/PA700, PA28αβ, PA28γ, PA200, and p97 (*14, 15, 36*). The best documented physiologic roles for proteasomes are mediated by these holoenzyme complexes, thereby raising questions about the physiologic need for inhibition of 20S proteasomes that are catalytically latent. PI31 might regulate cellular proteasome function by controlling the assembly of these holoenzymes. In fact, PI31 exerts such a function *in vitro* by physically blocking activator binding to α subunit rings (*21, 37*). We are unaware, however, of any evidence for such an effect in intact cells. PI31 has no direct effect on the activity of proteasome holoenzymes featuring activating regulators bound at each α ring. However, proteasomes composed of an activator complex on one end of 20S and PI31 on the other should feature three uninhibited catalytic subunits. The functional status of such a “hybrid” proteasome complex has not been determined experimentally and will require isolation of a homogenous enzyme population to document (*38*). Regardless of functional status, this type of PI31-containing complex appears to be responsible for the transport of proteasomes along axons to presynaptic terminals of motor neurons (*39*). In this context, PI31 functions as an adaptor that physically links the proteasome to motor proteins. The generality of this novel function for the transport of proteasomes to other intracellular locations and in other cell types remains to be determined. .

Finally, despite the lack of existing evidence, it is possible that PI31 exerts its established *in vitro* inhibitory function on cellular 20S proteasomes. There are many reports of roles for isolated 20S proteasomes in the cellular protein degradation (*40–42*). Although the mechanisms by which these normally latent enzymes are activated in the absence of regulatory proteins are not well understood, PI31 inhibition might serve a regulatory role to terminate their intended function. PI31 might also serve to inhibit 20S proteasomes that are inappropriately activated and require inhibition to avert cellular damage by unintended proteolysis. Additional work will be required to document such function for PI31 and to determine whether and why they might be different for 20Sc and 20Si.

## EXPERIMENTAL PROCEDURES

### Purification of constitutive and immuno-20S proteasomes

Catalytically latent constitutive (20Sc) and immuno-(20Si) 20S proteasomes were purified from bovine red blood cells and bovine spleen, respectively, as described previously (*13, 43*).

### Native polyacrylamide gel electrophoresis (PAGE)

**N**ative-PAGE was conducted as described previously using 3-8% acrylamide Tris-acetate gels (Invitrogen) (*44*). Samples were electrophoresed in using Tris-borate running buffer at 100 V for 3 hours at 4°C and either stained with Coomassie Blue R-250 or analyzed by zymography as described below.

### Expression and purification of recombinant PI31

Recombinant human PI31 (hPI31) was expressed and purified as described previously (*21*). In brief, PI31 cDNA was subcloned into a pET-28a(+) expression vector (Novagen) at the *Nde*I and *Hin*dIII restriction sites containing an amino-terminal polyhistidine sequence (His·Tag^TM^ [MGSSHHHHHHSSGLVPAGSH], Novagen) for affinity purification and a thrombin cleavage site for tag removal. The resulting cDNA sequence was verified. PI31 was expressed in *E. coli* BL21 Star (DE3) cells (Novagen) as described. Three-to-six hours after IPTG induction and incubation at 37°C, cells were harvested by centrifugation, lysed, and sonicated. A soluble extract was prepared by centrifugation at 14,000 × *g* for 30 min and applied to Ni-NTA His-bind Resin (Millipore Sigma) equilibrated with binding buffer (50 mM sodium phosphate, pH 7.6, 500 mM NaCl, 10 mM imidazole) and incubated for 1 hour. The resin was washed with 50 mM sodium phosphate, pH 7.6, 500 mM NaCl, and 20 mM imidazole. Bound protein was eluted with wash buffer containing 250 mM imidazole. The eluted protein was dialyzed extensively against Buffer H (20 mM Tris-HCl, pH 7.6, 20 mM NaCl, 1 mM EDTA, 5 mM β-mercaptoethanol) and when necessary purified further by ion exchange chromatography on DEAE Sephacyl using a 20-350 mM NaCl gradient in Buffer H. PI31 purity was assessed by SDS-PAGE and defined as a single band of approximately 31,000 Da upon staining with Coomassie Blue R-250. The identity of the band as PI31 was verified by western blotting with antibodies against the His-tag and PI31. Purified PI31 was concentrated to using Amicon XM10 membrane and stored at -80° C until use. As noted previously, we found that the use of thrombin to remove the His-tag typically cleaved PI31 at multiple points in the C-terminal domain. We therefore employed PI31 containing the N-terminal His-tag for these experiments. Previous work demonstrated that the His-tag had no demonstrable effect on PI31 function when compared to untagged native PI31 purified from bovine or human red blood cells (unpublished data and ref (*21*)).

### Expression and purification of recombinant PI31 mutants

Mutants of human PI31 were engineered and expressed in *E. coli* by PCR mutagenesis using wild-type PI31 cDNA, as described previously (*21, 35*). In brief, primers were designed to introduce the *Nde*I and *Hin*dIII restriction sites flanking the PI31 deletion sequences for subcloning into the pET-28a(+) expression vector at the *Nde*I and *Hin*dIII restriction sites. The sequences of all mutant constructs were verified. Expression of mutant PI31 proteins in *E. coli* and protein purification were performed as described above for wild-type PI31.

### Proteasome activity assays

Proteasome activity was assessed by the following assays.

#### Hydrolysis of 7-amino-4-methylcoumarin (AMC)-linked peptide

Proteasome activity was measured by determining rates of hydrolysis of AMC-linked peptides such as Suc-LLVY-AMC, essentially as described previously (*44*). In brief, 20S proteasomes and any other components of specific reactions, such as PI31 and activator peptides or proteins were preincubated for 10 mins in a buffer of 50 mM Tris-HCl, pH 7.6 at 37°C in a final volume of 50 μl. Reactions were initiated by addition of 150 μl Suc-LLVY-AMC buffered with 20 mM Tris-HCl, pH 7.6 and 20 mM NaCl; final substrate concentration was 50 μM. Rates of production of free AMC were determined by continuous monitoring (one reading/min for 20 mins) of fluorescence in a BioTek Synergy plate reader (380 nm Ex, 460 nm Em). Background fluorescence from reactions containing no enzyme was subtracted from enzyme-containing samples. In given experiments, reactions were conducted in triplicate and rates of hydrolysis were expressed as mean values of arbitrary fluorescent units (AFU)/min/μg protein, ± standard deviation.

#### Hydrolysis of fluorescein isothiocyanate (FITC)-casein

Proteasome activity was measured by determining rates of hydrolysis of fluorescein isothiocyanate (FITC)-labeled casein, essentially as described previously (*45*). In brief, α-casein (500 mg in 50 ml of 50 mM sodium carbonate buffer, pH 9.5) was incubated with 50 mg FITC for 8 hours at room temperature. The solution was then dialyzed extensively against 25 mM Tris-HCl, pH 7.5. The dialyzed protein was diluted to 2 mg/ml and stored at – 20°C. Rates of proteasome-catalyzed degradation of FITC-casein were measured by production of tricholoracetic acid-soluble FITC peptides during the incubation. FITC-casein (5 μg/assay) was incubated in the presence or absence of 20S proteasome and any other reaction components as specified in individual figure legends. Assays were performed at 37°C in 50 mM Tris-HCl buffer, pH 7.6, and 20 mM NaCl, in a final volume of 50 μl. At various times of incubation reactions were terminated with trichloroacetic acid (TCA) to a final concentration of 10%. 1 mg/ml BSA was included as a carrier protein for complete precipitation. After 1 hr at 4°C, samples were centrifuged to pellet TCA-precipitated protein. 20 μl aliquots of the supernatant were added to 200 μl of 1 M Tris-HCl, pH 8.5. Fluorescence was quantitated using a BioTek Synergy plate reader at Ex 495 nm, Em 525 nm. Fluorescent values of samples incubated in the absence of proteasome and any other assay components were subtracted from corresponding values of sample incubated with proteasome. Under conditions of maximal proteasome activity, the production of TCA-soluble products was linear for up to 3 hours. Assays were conducted in triplicate and rates of hydrolysis were expressed as mean values of arbitrary fluorescent units (AFU)/min/μg protein, ± standard deviation.

#### Hydrolysis of α-synuclein

Proteasome activity was measured by determining rates of hydrolysis of α-synuclein. Recombinant α-synuclein containing a C-terminal His-tag was expressed in *E. coli* and purified as described previously (*46*). Proteasome-dependent degradation of α-synuclein was determined by measuring the rate of disappearance of intact α-synuclein-His on SDS-PAGE. 20S proteasomes (200 nM) were incubated with α-synuclein in the presence or absence of PI31 (1 μM, or as specified for given experiments) and in the presence or absence of 5 mM z-YA, in 50 mM Tris-HCl, pH 7.6 and a final volume of 50 μl. Reactions were terminated by addition of SDS sample buffer. Samples were subjected to SDS-PAGE and stained with Coomassie Blue R250. α-synuclein content was quantified using Image Studio software (LiCOR).

#### Proteasome activity-based probe labelling

Latent, constitutive and immuno-proteasome (131 nM) was incubated with 5 mM zYA or buffer controls for 5 minutes at 37°C in 50 mM Tris-HCl (pH 7.6). Recombinant His-PI31 (13.1 µM) or buffer control was then added for 10 minutes at 37°C to allow for binding to 20S core particles. The proteasome activity-based probe Me_4_BodipyFL-Ahx_3_Leu_3_-VS covalently labels all catalytic subunits of constitutive and immuno-proteasomes (*47*). Proteasomes were labeled with 500 nM activity probe for 30 minutes at 37°C. The reaction was terminated with SDS sample buffer. Proteasome subunits (1.5 µg total proteasome) were separated by 12% SDS-PAGE and transferred to nitrocellulose membranes. Proteasome subunit labeling was visualized on an Odyssey M imager (LiCOR) at Ex 520 nm, Em 590 nm and quantified in Image Studio (LiCOR).

#### Zymography with native PAGE

Zymography of proteasomes after native PAGE was conducted as described previsously (*44*). In brief, after electrophoresis, gels were were soaked under 50 µM Suc-LLVY-amc in 20 mM Tris-HCl (pH 7.6) for 30 minutes at 37°C with or without 0.03% SDS or 5 mM z-YA activator, as described in figure legends, and imaged under trans-illumination with a ChemiDoc MP imager (BioRad) using the fluorescein channel.

### Mass spectrometry

Immuno-proteasome (2.2 nM) was activated with 5 mM z-YA at 37°C for 5 minutes. His-PI31 (5.6 nM) was then added to activated 20Si and allowed to degrade for 90 minutes at 37°C in 50 mM Tris-HCl (pH 7.6). Binding buffer was added to the sample to a final concentration of 50 mM sodium phosphate (pH 7.4), 500 mM NaCl, 10 mM imidazole, and the sample was incubated with 50 µl Ni-NTA resin (Novagen) at 4°C for 30 minutes with gentle shaking. HisPI31 degradation product was purified after 4 washes with binding buffer containing 20 mM imidazole, and eluted with binding buffer containing 250 mM imidazole. As a control, HisPI31 was purified in the same manner without incubation with 20Si. Intact protein samples were analyzed by LC/MS, using a Sciex X500B QTOF mass spectrometer coupled to an Agilent 1290 Infinity II HPLC. Samples were injected onto a POROS R1 reverse-phase column (2.1 x 30 mm, 20 µm particle size, 4000 Å pore size) and desalted. The mobile phase flow rate was 300 μl/min and the gradient was as follows: 0-3 min: 0% B, 3-4 min: 0-15% B, 4-16 min: 15-55% B, 16-16.1 min: 55-80% B, 16.1-18 min: 80% B. The column was then re-equilibrated at initial conditions prior to the subsequent injection. Buffer A contained 0.1% formic acid in water and buffer B contained 0.1% formic acid in acetonitrile. The mass spectrometer was controlled by Sciex OS v.3.0 using the following settings: Ion source gas 1 30 psi, ion source gas 2 30 psi, curtain gas 35, CAD gas 7, temperature 300 °C, spray voltage 5500 V, declustering potential 125 V, collision energy 10 V. Data were acquired from 400-2000 Da with a 0.5 s accumulation time and 4 time bins summed.

### Glycerol density gradient centrifugation of PI31-20S complexes

Purified constitutive and immuno-proteasome 20S core particles (1.25 µM) were pre-incubated with or without 5 mM z-YA in 20 mM Tris-HCl (pH 7.6) at 37°C for 5 minutes. Recombinant His-PI31 was then added at a 2:1 molar ratio (2.5 µM) and further incubated for 10 minutes at 37°C. In cases where constitutive and immuno-proteasome irreversible inhibitors were used, 1.8 µM epoxomicin (for 20Sc) and 430 µM ONX-0914 (for 20Si) were added prior to incubation. Following the pre-incubation, samples were subjected to glycerol density gradient centrifugation (10-40% glycerol) was conducted as described previously in an Optima TL ultracentrifuge, Beckman Coulter (*35*). Following centrifugation, 100 µl fractions were collected and denatured in SDS sample buffer and subjected to SDS-PAGE and western blotting as indicated in specific figures. Proteins detected in the first 13 fractions (10-30% glycerol).

### Western blotting and antibodies

Western blotting was performed as previously described (*35*). Proteins transferred to nitrocellulose membranes were blocked and blotted using primary antibodies against: 6xHis (mouse monoclonal, Invitrogen MA1-135); 20S α4 PSMA7 (mouse monoclonal anti-human, Enzo PW8120); PI31 C-terminus residues 241-271 (rabbit polyclonal anti-human, Enzo BML-PW9710); 20S β2i PSMB10 (rabbit polyclonal anti-human, Cell Signaling 78385S); Rabbit polycolonal antibodies against human 20S β1C PSMB6, β1i PSMB9, β5c PSMB5, and β5i PSMB8 subunits were prepared in our laboratory using C-terminal peptides of the respective subunits, as described and characterized previously (*48*). All primary antibodies were typically used at 1:1000-5000 dilution. Secondary antibodies IRDye 800CW Goat anti-mouse IgG (P/N 926-32210) and IRDye 680RD Goat anti-rabbit IgG (P/N 926-68071) were obtained from LiCOR and used at 1:10,000 dilution. Proteins were scanned using an Odyssey M imager and analyzed in Empiria Studio or Image Studio software (LiCOR).

### Statistical analysis

Proteasome activities between the two 20S isoforms under different treatments (e.g. z-YA activation or PI31 inhibition) were analyzed by ANOVA with the indicated amount of factors (2-way or 3-way ANOVA) for specific experiments, as described in figure legends. Post-hoc pair-wise comparisons were made using Tukey’s HSD test where appropriate. For analyses of activity across PI31 concentration ranges, repeated measures 2-way ANOVA was used for 20S isoform as a factor repeated across PI31 concentrations. Differences in activity at PI31 concentration-matched treatments were analyzed by Šídák’s multiple comparisons test. Statistics were calculated using GraphPad Prism (version 10.1.1)

## Acknowledgements

This work was supported by a grant from the National Institutes of Health (GM129088 to GND). We thank Dr. Andrew Lemoff of the UT Southwestern Proteomics Core Facility for performing intact protein mass spectrometry.

## Author contributions

JW, AK, and GND designed the research. JW and AK performed the experiments. GND and JW wrote the manuscript. All authors provided data analysis and input and editing of the final manuscript version.

## Competing interests

The authors declare no competing interests.

## Notes

### Competing Interest Statement

The authors have declared no competing interest.

